# Position-dependent effects of cytosine methylation on *FWA* expression in *Arabidopsis thaliana*

**DOI:** 10.1101/774281

**Authors:** Thanvi Srikant, Anjar Wibowo, Rebecca Schwab, Detlef Weigel

**Affiliations:** Max Planck Institute for Developmental Biology, Max-Planck-Ring 9, 72076 Tübingen, Germany; Faculty of Science and Technology, Airlangga University, Kampus C, Mulyorejo, Surabaya City, East Java 60115, Indonesia

## Abstract

Gene expression can be modulated by epigenetic modifications to chromatin, and variants of the same locus distinguished by fixed, heritable epigenetic differences are known as epialleles. DNA methylation at cytosines is a prominent epigenetic modification, particularly in plant genomes, that can modulate gene expression. There are several examples where epialleles are associated with differentially methylated regions that affect the expression of overlapping or close-by genes. However, there are also many differentially methylated regions that have not been assigned a biological function despite their proximity to genes. We investigated the positional importance of DNA methylation at the *FWA (FLOWERING WAGENINGEN)* locus in *Arabidopsis thaliana*, a paradigm for stable epialleles. We show that cytosine methylation can be established not only over the well-characterized *SINE-*derived repeat elements that overlap with the transcription start site, but also in more distal promoter regions. *FWA* silencing, however, is most effective when methylation covers the transcription start site.

## INTRODUCTION

Methylation of cytosine nucleotides in DNA is a prominent epigenetic mark in plant and animal genomes (1). It is found mostly over transposons and repeat elements, consistent with its primary function in silencing their transcriptional activity in association with methylation at lysine 9 of histone 3 (H3K9) (2). Mutants with defects in *METHYLTRANSFERASE 1* (*MET1*), encoding the major methyltransferase maintaining cytosine methylation in *Arabidopsis thaliana*, display various phenotypic abnormalities such as delayed flowering, dwarfism, and sterility, with increasing severity during successive rounds of inbreeding (3, 4). Genomes of *met1* mutants are largely hypomethylated at CG dinucleotides that usually inherit cytosine methylation faithfully during DNA replication through MET1 activity (5). Phenotypic abnormalities do, however, not require whole-genome changes in cytosine methylation, as a recent study describes how hypomethylation at a few select loci is sufficient to establish quantitative resistance to a pathogenic oomycete, *Hyaloperonospora arabidopsidis (Hpa)* (6).

Transposable elements and repeats in the *A. thaliana* genome are mostly confined to heterochromatic regions, such as pericentromeres and telomeres. Others are distributed in euchromatic regions, and their proximity to protein-coding genes has been associated with constitutive or induced silencing of these proximal genes upon changes in cytosine methylation (7, 8). There are several examples of loci with alternative states of cytosine methylation and associated gene expression; the variants are known as epialleles. One example comes from the *SDC* (*SUPPRESSOR OF ddc*) locus with a direct tandem repeat in its promoter. When methylation at the repeat is absent, *SDC* is expressed, resulting in a dwarfed phenotype (9). Epialleles can also form following stress exposure; treatment with the 22-amino-acid peptide flg22, an immune-response inducing fragment of the bacterial flagellin protein, causes differential methylation of helitron-derived repeats lying within a 3 kb promoter region of the defense gene *RESISTANCE METHYLATED GENE 1* (*RMG1*), and ensuing activation of *RMG1* expression (10). Furthermore, cytosine methylation has been shown to modify gene expression when located farther away from the gene body, such as in the case of *FLOWERING LOCUS T* (*FT*), where methylation on two enhancers located 5 kb upstream and 1 kb downstream of the gene can repress transcriptional activity (11). This was shown by experimentally targeting cytosine methylation to these enhancers using Inverted Repeat-Hairpins (IR-Hairpins), which lead to the downregulation of *FT* expression and delayed flowering.

While these examples provide substantial evidence for methylation-dependent transcriptional changes, the requirements for cytosine methylation to exert this function is not well understood, as not all cytosine methylation, even when densely focused in methylated regions, triggers silencing of adjacent genes (12).

To address such functional differences, we investigated the promoter of the well-characterized *Arabidopsis thaliana fwa*-1 epiallele. The *FWA* (*FLOWERING WAGENINGEN*) locus (At4G25530) harbors two sets of tandem repeats originating from a *SINE3* retrotransposon (13). These repeats overlap the promoter and the *FWA* transcribed sequence, and are covered by dense CG methylation in wild-type plants. Throughout vegetative development, *FWA* is transcriptionally inactive, and activated only in the female gametophyte and endosperm by maternal imprinting, when DNA methylation is erased (14). Methylation at the repeats is also absent in *fwa*-1 epimutants, where the gene is constitutively active, which results in late flowering (15–18).

It is known that the presence of cytosine methylation at a specific position, the *SINE*-derived repeat elements, imposes transcriptional silencing on the entire locus. We made use of the unmethylated promoter in the *fwa*-1 epiallele and asked the reverse, whether *FWA* silencing can be triggered only by cytosine methylation at these repeats, or whether it can be similarly induced when cytosine methylation is artificially directed to other, non-repetitive, promoter elements.

## RESULTS

### Hairpins directing methylation to the *FWA* promoter

We chose three regions of 100 or 200 basepairs (bp) in length, located approximately 100, 500 and 700 bp upstream of the *FWA* transcription start site (Table 1 and Figure 1), and generated inverted repeat (IR)-hairpins (19), intended to introduce cytosine methylation at these regions that are otherwise unmethylated in *fwa-1* mutants.

**Table 1:** IR-Hairpin constructs targeting the *FWA* promoter

| Construct | Length (bp) | Orientation of 5' arm target sequence relative to *FWA* transcription | Distance from *FWA* transcription start site |
| --- | --- | --- | --- |
| 1A | 100 | sense | -122 to -22<br>(Overlap with short tandem repeats) |
| 1B | 100 | antisense | -122 to -22<br>(Overlap with short tandem repeats) |
| 2A | 100 | sense | -659 to -559 |
| 2B | 100 | antisense | -659 to -559 |
| 3A | 200 | sense | -936 to -736 |
| 3B | 200 | antisense | -936 to -736 |

**Figure 1.**
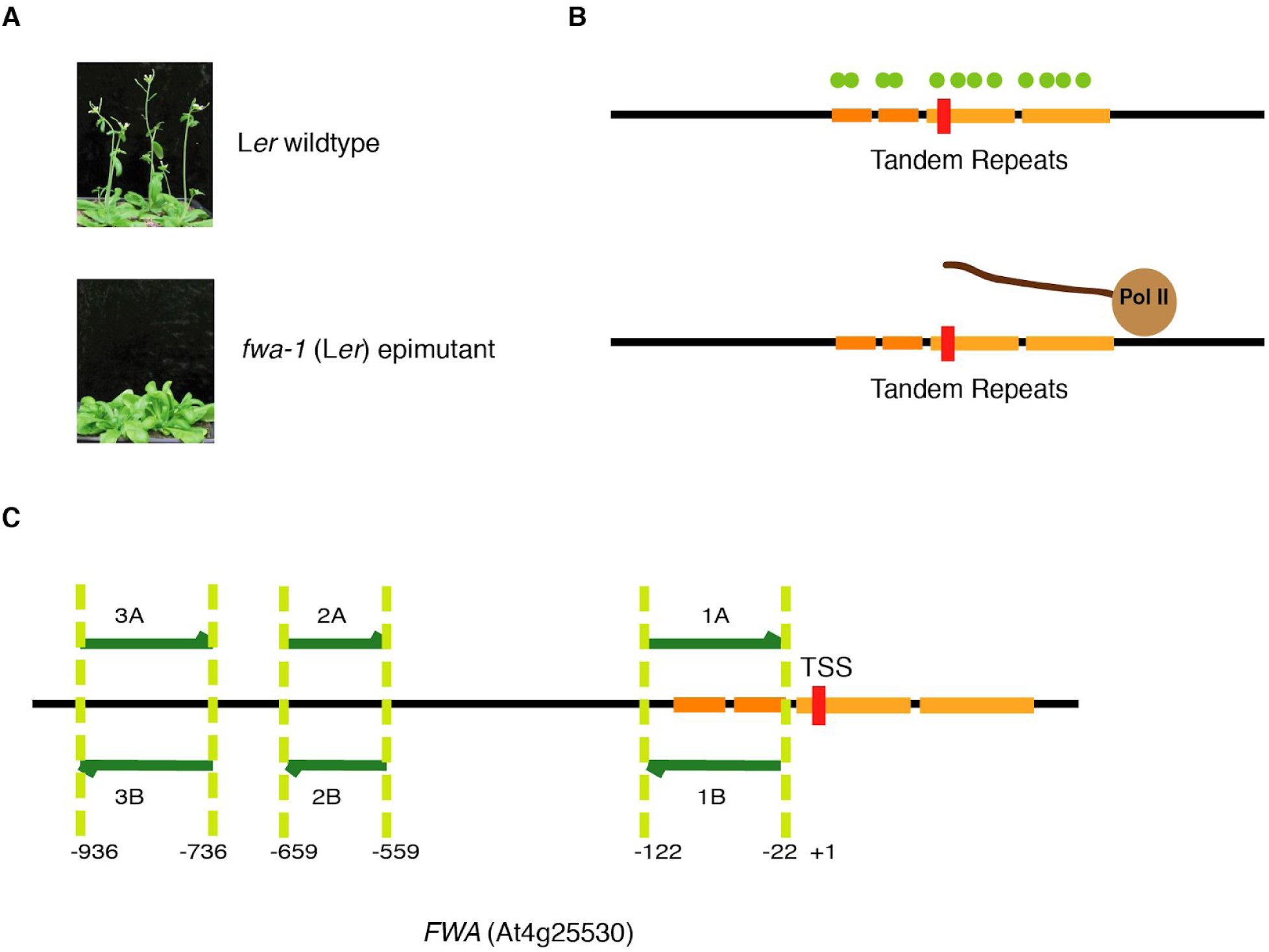
Methylation tiling in the *FWA* promoter. The diagram illustrates the *FWA* locus and the regions targeted for methylation tiling. (A) Late flowering phenotype of *fwa-1* (B) Release of methylation in the tandem repeats which overlap the TSS activates the *FWA* transcript (C) Three Regions in the *FWA* promoter chosen for hairpin-induced targeted methylation. Red line indicates the transcription start site (TSS) and the orange horizontal bars indicate the tandem repeats.

For each region, we generated two IR-hairpins, differing in the orientation of sense and antisense sequences within the hairpin (Table 1, Figure 1, Methods).

### Region-specific methylation effects on *FWA* activity

We transformed late-flowering *fwa-1* mutants with IR-hairpin transgenes as described above (Table 1), and identified homozygous insertion lines to monitor flowering time as a proxy for *FWA* activity. Corroborating previous work (15–18, 20), hairpins derived from sequences overlapping the repeat elements surrounding the transcription start site of *FWA* (lines 1A and 1B) accelerated flowering. This was different for hairpins derived from more distal promoter sequences (lines 2A, 2B, 3A and 3B), where we did not observe obvious flowering differences compared to parental *fwa-1* individuals (Figure 2 and Figure 3).

**Figure 2.**
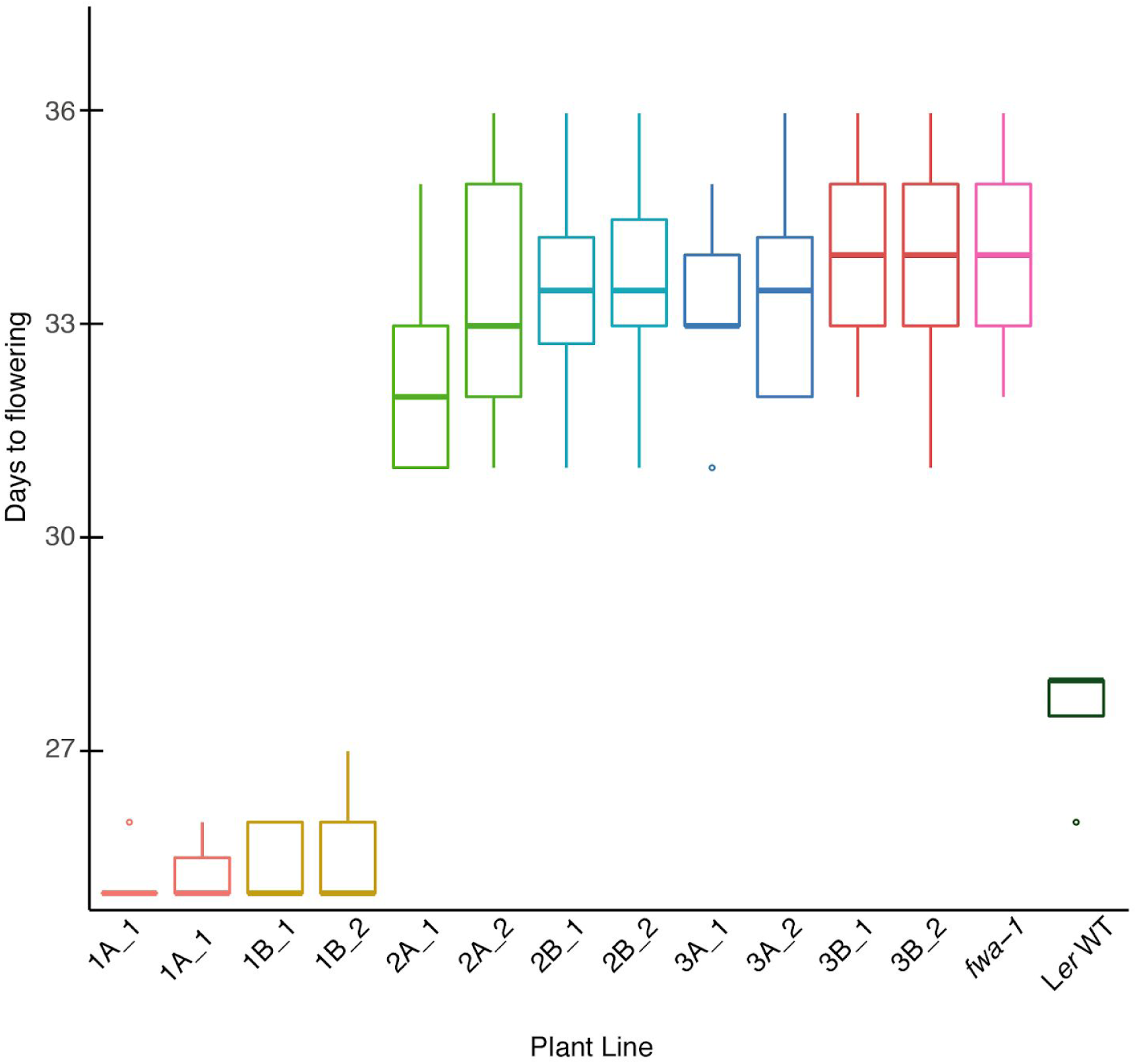
Flowering time in various transgenic lines. Box plots showing days to flowering in L*er* WT, *fwa-1* and various transgenic lines in the *fwa*-1 background (with 10 randomized replicates for each line).

**Figure 3.**
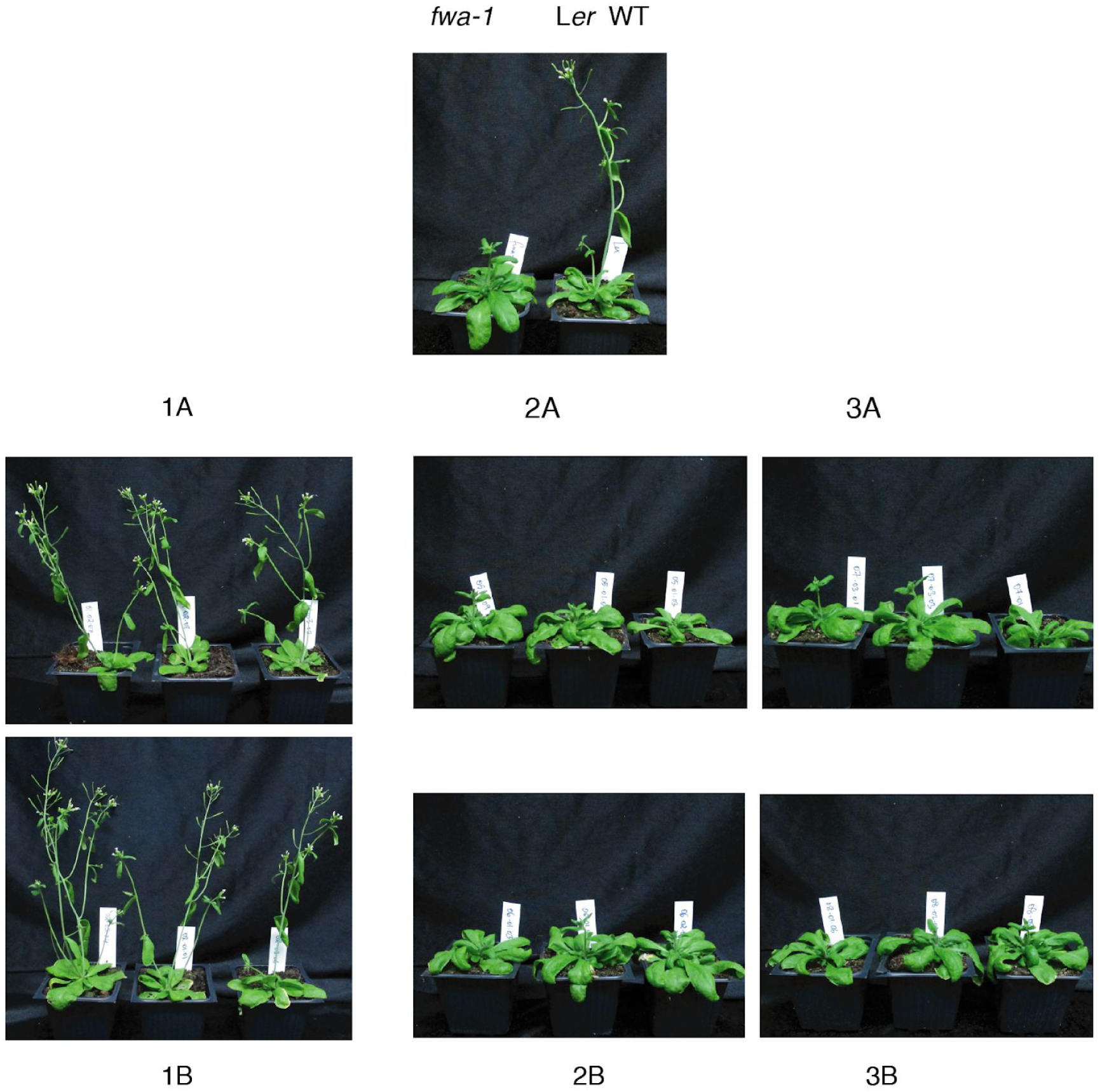
Phenotypes of transgenic lines. Transgenic lines and control plants at 28 days after germination.

Consistent with these observations, *FWA* transcripts in lines 1A and 1B accumulated to levels much lower than in parental *fwa*-1, and more similar to those in early-flowering L*er* wild-type plants, indicative of effective gene silencing (Figure 4). Similarly, lines in which flowering time was unaltered showed less reduction (lines 2A, 2B, 3A) or no reduction (line 3B) of *FWA* mRNA accumulation (Figure 4).

**Figure 4.**
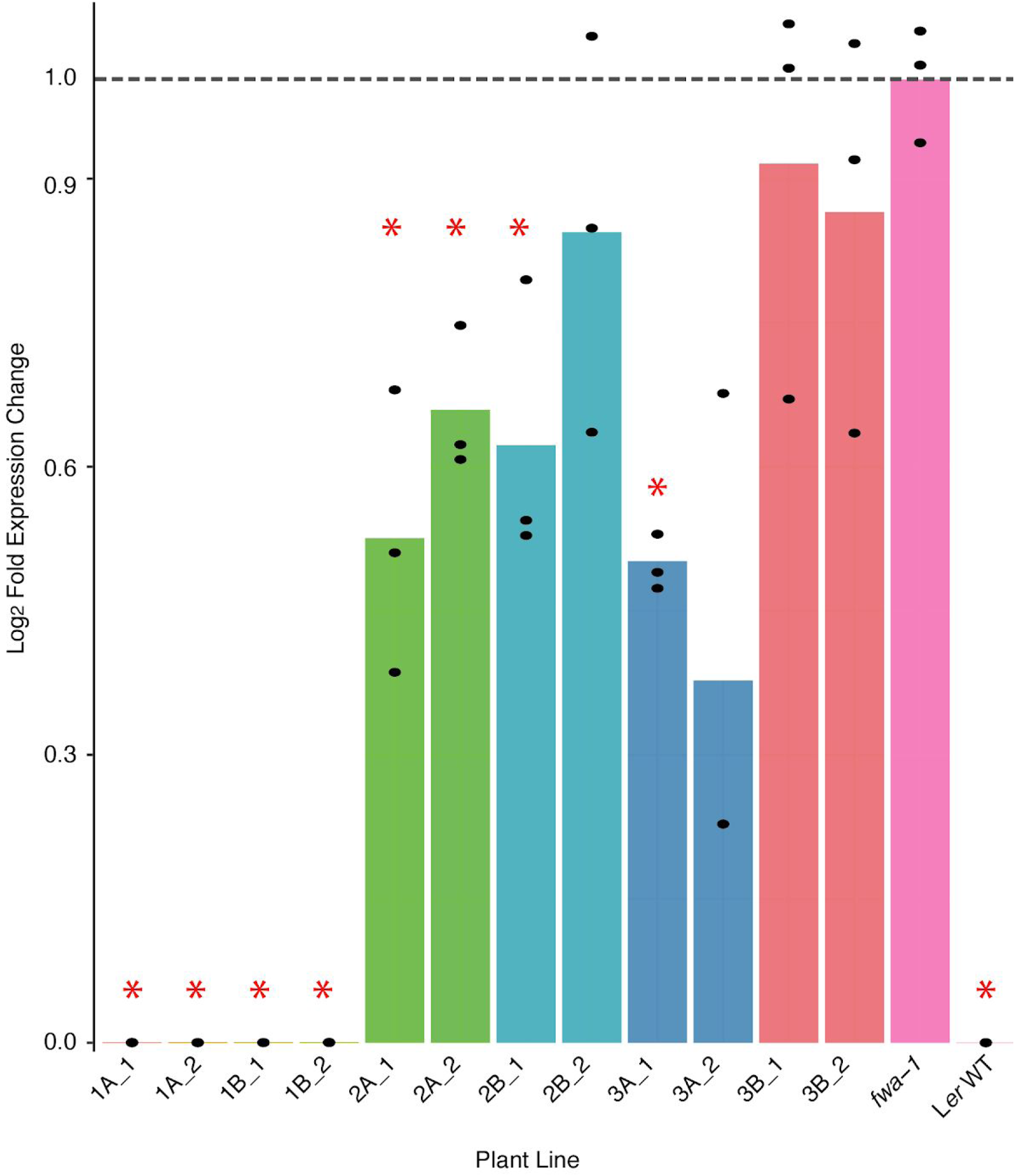
*FWA* mRNA accumulation in plants expressing IR-hairpins. Expression of *FWA* in L*er* WT, *fwa-1* and various transgenic lines carrying the hairpin-construct in sense and antisense orientations. Gene expression was measured by RT-qPCR. Bars show mean log_2_-fold change compared to *fwa-1*, with black dots representing pooled biological replicates. The red asterisks represent statistical significance (p<0.05 for a *Student’s t-test*).

### Establishment of cytosine methylation at distal promoter regions

As IR-hairpins derived from more distal regions of the *FWA* promoter did not induce sufficient *FWA* silencing to alter the timing of flowering, we asked whether cytosine methylation had been effectively established at the targeted sequences using whole-genome bisulfite sequencing. As shown in Figure 5, all six transgenes efficiently introduced methylation in the targeted regions to comparable levels. The effect of the transgene on methylation was stable over at least three generations, and indicates that cytosine methylation at this locus can be introduced not only at the *SINE*-derived repeat elements. We further conclude that methylation at more distal promoter elements was less effective in *FWA* silencing, which may be due to the distance from the transcription start site.

**Figure 5.**
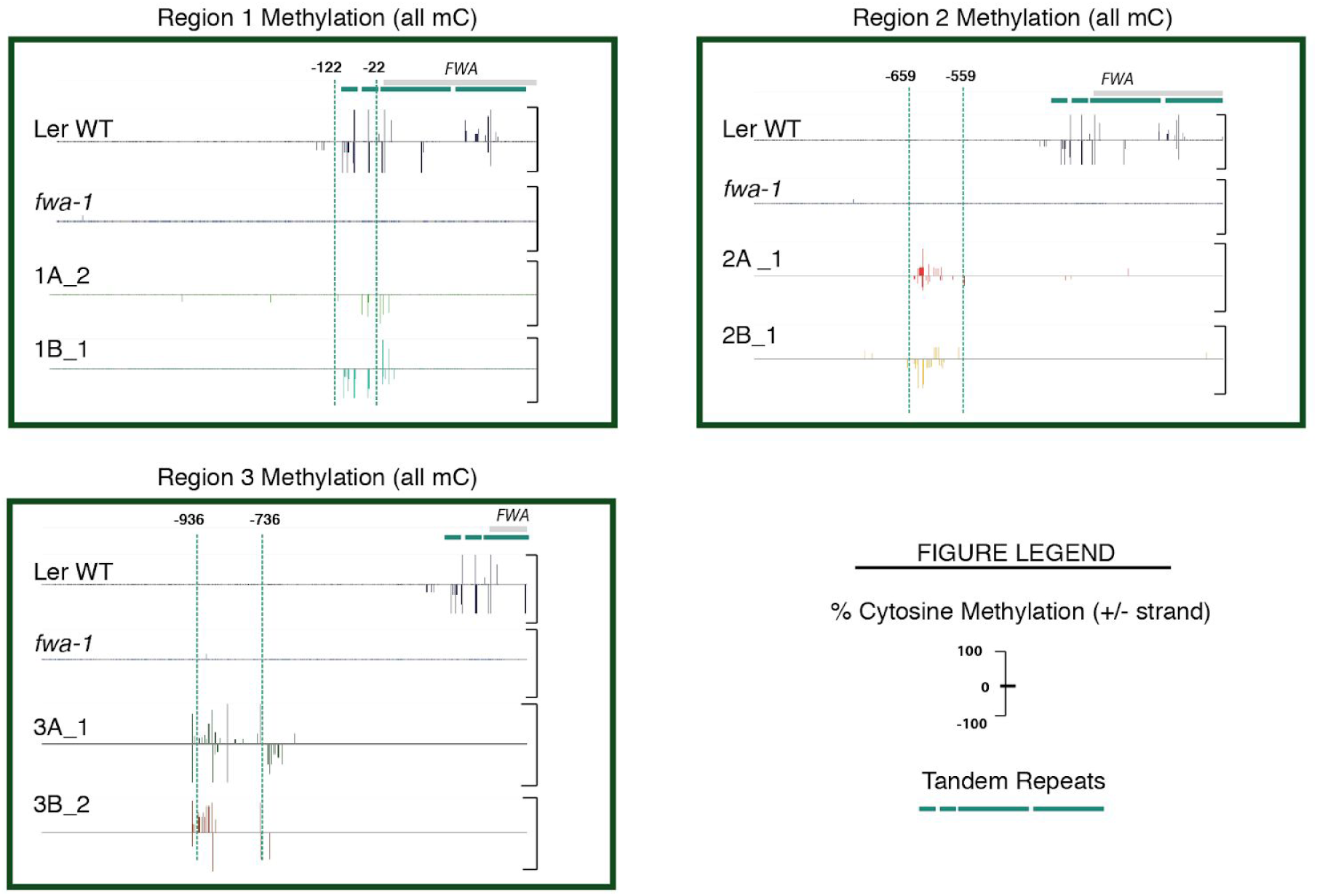
Methylation profile of targeted regions in transgenic lines. Hairpin-induced methylation for various transgenic lines. Each panel shows the methylation status of the targeted region in the L*er* WT, *fwa-1* and transgenic lines carrying the hairpin-construct in sense and antisense orientations. Green dotted lines demarcate the positions of each region. The vertical lines represent percentage methylation at every cytosine in both strands.

## DISCUSSION

Cytosine methylation can be altered at many genomic regions under environmental stress. For example, phosphate starvation in rice and *Arabidopsis thaliana* results in the generation of hypermethylated differentially methylated regions (DMRs), many of which are found over transposable elements that are close to genes induced upon such nutrient starvation (21). On the contrary, some studies report that stress-induced methylation and gene expression are not necessarily correlated with one another. In *A. thaliana*, most genomic regions exhibiting differential methylation under drought are not associated with expression changes of adjacent genes, with only 2 of 468 drought-response genes being linked to drought DMRs (22). These studies support the observation that methylation changes often flank protein-coding genes, yet only a few of them have been associated with a biological function. Several epialleles characterized in *A. thaliana* are marked by differential cytosine methylation in transposon-derived repetitive elements located proximal to the transcription start site (for example, *SDC*, (9); *RMG1*, (10); *HDG3*, (23)). This pattern also holds true for the *FWA* locus, where *SINE*-derived tandem repeats that overlap with the transcription start site are heavily methylated and silence the gene.

Transcription is initiated in the proximity of DNA elements providing a platform for the recruitment and assembly of polymerase II-containing complexes. Their binding is modulated by enhancers and repressors, which themselves may recognize distal sequence elements. It is likely that cytosine methylation over elements of polymerase docking may directly or indirectly hinder effective protein recruitment. When located in regions distal to the transcription start site, methylation-based silencing may be achieved by the inhibition of transcriptional enhancers (24), and by altering short or long-distance chromatin interactions (25), thus affecting accessibility to the transcription machinery. Such a mechanism may possibly account for our observations, that methylation targeted in Regions 2 and 3, located more than 500 bp upstream of the transcription start site, only moderately impacts *FWA* transcript accumulation.

On examining publicly available data from the PlantDHS Browser (26), we observed that the 1-kb promoter region immediately upstream of the *FWA* transcription start, including all three regions targeted for methylation, has a uniform state of chromatin accessibility (Figure 6), not only when in the silent wild-type state, but also in the unsilenced state in *ddm*-1 mutants, which express *FWA* due to hypomethylation of the tandem repeats (27). Therefore, it may be expected that one would see a similar transcriptional readout upon modulating different regions within the same inaccessible sequence. At least in the *FWA* locus, this is not the case. Our results indicate that, while IR hairpin-induced methylation can be successfully introduced in the distal *FWA* promoter, this methylation can induce only moderate downregulation of *FWA* transcription, in contrast to very potent effects when methylating elements more proximal to the transcription start site.

**Figure 6.**
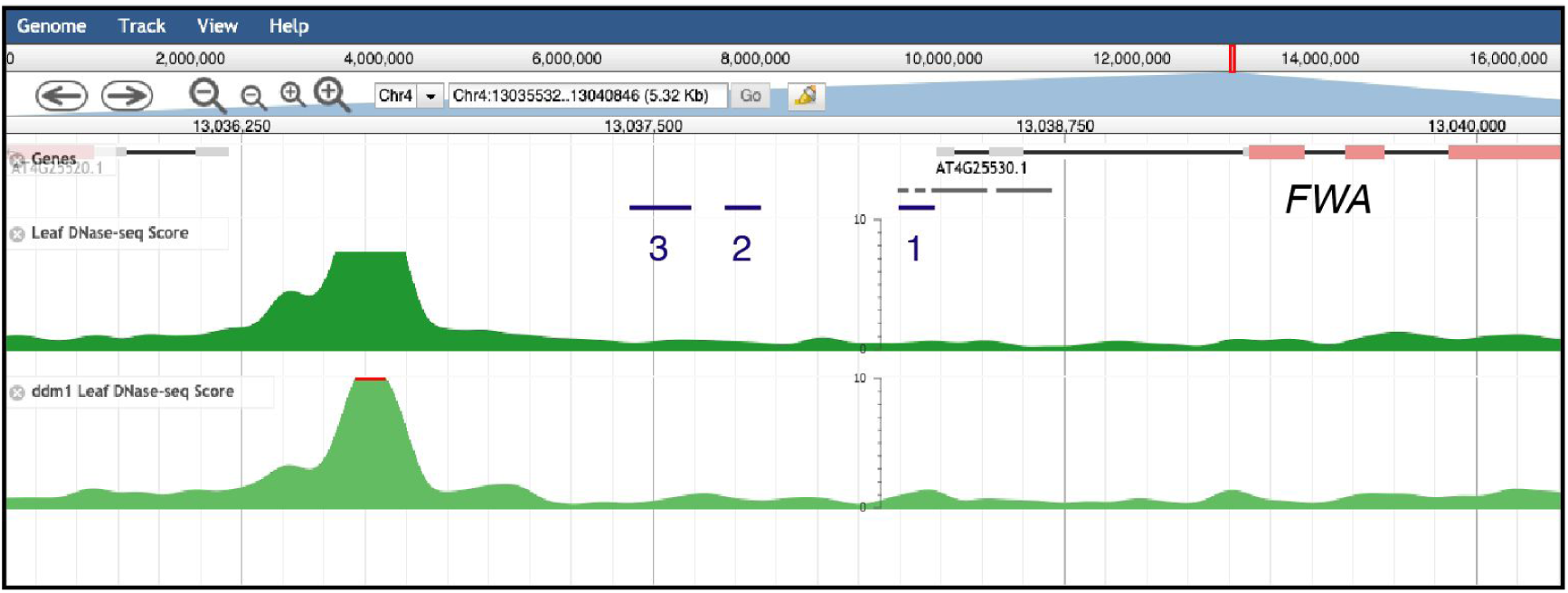
Chromatin conformation at the *FWA* locus. Screenshot from the PlantDHS browser (http://plantdhs.org/) showing the DNaseI-seq profile of the *FWA* locus in leaf tissue of Col-0 wild-type (top panel) and *ddm-1* mutant (bottom panel) backgrounds. Grey bars below the *FWA* gene annotation indicate the tandem repeats; blue bars indicate the three regions targeted for methylation in this study.

It will be interesting to examine the chromatin state and conformation of the *FWA* promoter in our transgenic lines, to understand whether they can explain the differential transcriptional downregulation that we observed.

This study was focused on the *FWA* locus; further investigation into differential promoter methylation at other genomic loci is required to dissect the mechanism behind methylation-dependent control on downstream transcription processes, and ultimately uncover its adaptive and biological function.

## METHODS

### Cloning of inverted-repeat hairpin (IR-Hairpin) constructs

Three 100 bp – 200 bp regions in the *FWA* promoter were chosen for IR-hairpin construction (Table 1 and Supplementary Table S1), and cloned in two ways: All ‘A’ constructs carry sequences in sense orientation relative to the open-reading frame of the *FWA* locus in the hairpin 5’ arm, while the ‘B’ constructs are reversed.

Hairpins, including flanking attB gateway sites were synthesised (GeneArt), PCR-amplified and transferred to pDONR207 (Invitrogen) using BP Clonase II. Recombinant entry clones were subjected to LR Clonase II reaction with the pJawohl-ACT2 destination vector (28) and introduced into *E.coli* DH5α (Invitrogen) cells by heat-shock transformation. Colonies carrying the hairpin construct in the correct orientation were verified by Sanger sequencing (oligonucleotide sequence provided in Supplementary Table S1).

Recombinant plasmids (listed in Supplementary Table S4) were introduced into *Agrobacterium tumefaciens* strain GV3101(pMP90RK) (29) by electro-transformation and grown in selective LB medium. *Arabidopsis thaliana fwa-1* mutants (16) were transformed with a floral dip protocol (30). Transgenic seeds were selected with 1% BASTA (Sigma-Aldrich).

### RT-qPCR

Homozygous third-generation transformants (T_3_) derived from three independent parent lines were sown on half-strength MS (Duchefa Biochemie) plates with sucrose and grown under long day (16h light/8h dark) conditions at 23 °C for 10 days in a Percival chamber (Model CU −36L5, CLF Plant Climatics GmbH, Germany). For each line, 3 biological replicates of 20 pooled seedlings were subjected to RNA extraction (based on the LogSpin method, (31)). cDNA synthesis was carried out with an equimolar concentration of an Oligo(dT)18 primer and an *FWA* gene-specific primer (Supplementary Table S2) using the RevertAid First Strand cDNA synthesis kit (Thermo Fisher Scientific), followed by qPCR of a 120bp region within the *FWA* gene body (primer sequences provided in Supplementary Table S2). The housekeeping gene *ACTIN2* (*At3G18780*) was used as a control gene for the experiment (primer sequences provided in Supplementary Table S2).

### Plant growth and flowering time analyses

Seeds were sterilised by treatment with chlorine gas for 4 hours, followed by stratification in the dark at 4**°**C for 2 days in 0.1% agar. All plants were grown in controlled growth chambers at 23 **°**C, long day conditions (16h light/8h dark) with 65% relative humidity under 110 to 140 μmol m^−2^ s^−1^ light provided by Philips GreenPower TLED modules (Philips Lighting GmbH, Hamburg, Germany) with a mixture of 2:1 DR/W LB (deep red/white mixture with ca. 15% blue) and W HB (white with ca. 25% blue), respectively and watered at 2-day intervals.

Ten plants belonging to each transgene were grown on soil in a randomized–block design, to reduce position effects in the growth chamber. Flowering time was recorded when the primary inflorescence meristem was approximately 1cm in height.

### Bisulfite library preparation and sequencing

Genomic DNA was isolated using the DNeasy Plant Mini Kit (Qiagen) from pools of twenty 10-day old seedlings grown on the same MS-Agar plates as seedlings used for RT-qPCR. 100ng of genomic DNA was used to prepare Bisulfite libraries with the TruSeq Nano kit (Illumina, San Diego, CA, USA) according to the manufacturer’s instructions, with the modifications used in (12). The libraries were sequenced with a paired-end mode, at 125 million reads/library using an Illumina HiSeq3000 instrument.

### Processing of sequenced bisulfite libraries

Raw sequencing reads were aligned using Bismark (default parameters; (32) and mapped to the *A. thaliana* (Landsberg *erecta*) reference genome. Bam files generated after deduplication of reads were processed for identifying methylated cytosines using pipelines as previously described in (33). Methylated cytosines in the *FWA* locus were loaded onto the EPIC-CoGe browser (34) for visualisation.

## Author contributions

Study design: T.S., A.W., D.W. Experimental work and data analysis: T.S. Data interpretation: T.S., A.W., R.S., D.W. Drafting of initial manuscript: T.S. Editing and finalizing of manuscript: T.S., A.W., R.S., D.W.

## Acknowledgments

We thank Claude Becker (Gregor Mendel Institute, Vienna) for providing access to the pipeline for DMR calling, and Isaac Rodriguez (Gregor Mendel Institute, Vienna) for his expertise. The pJawohl-ACT2 and pDONR207 plasmid vectors were kindly shared by José Gutierrez-Marcos (University of Warwick). This study was supported by Marie Sklodowska-Curie Fellowship 751204-H2020-MSCA-IF-2016 (A.W.), DFG (ERA-CAPS AUREATE), and the Max Planck Society.

## SUPPLEMENTARY DATA

**Table S1.**
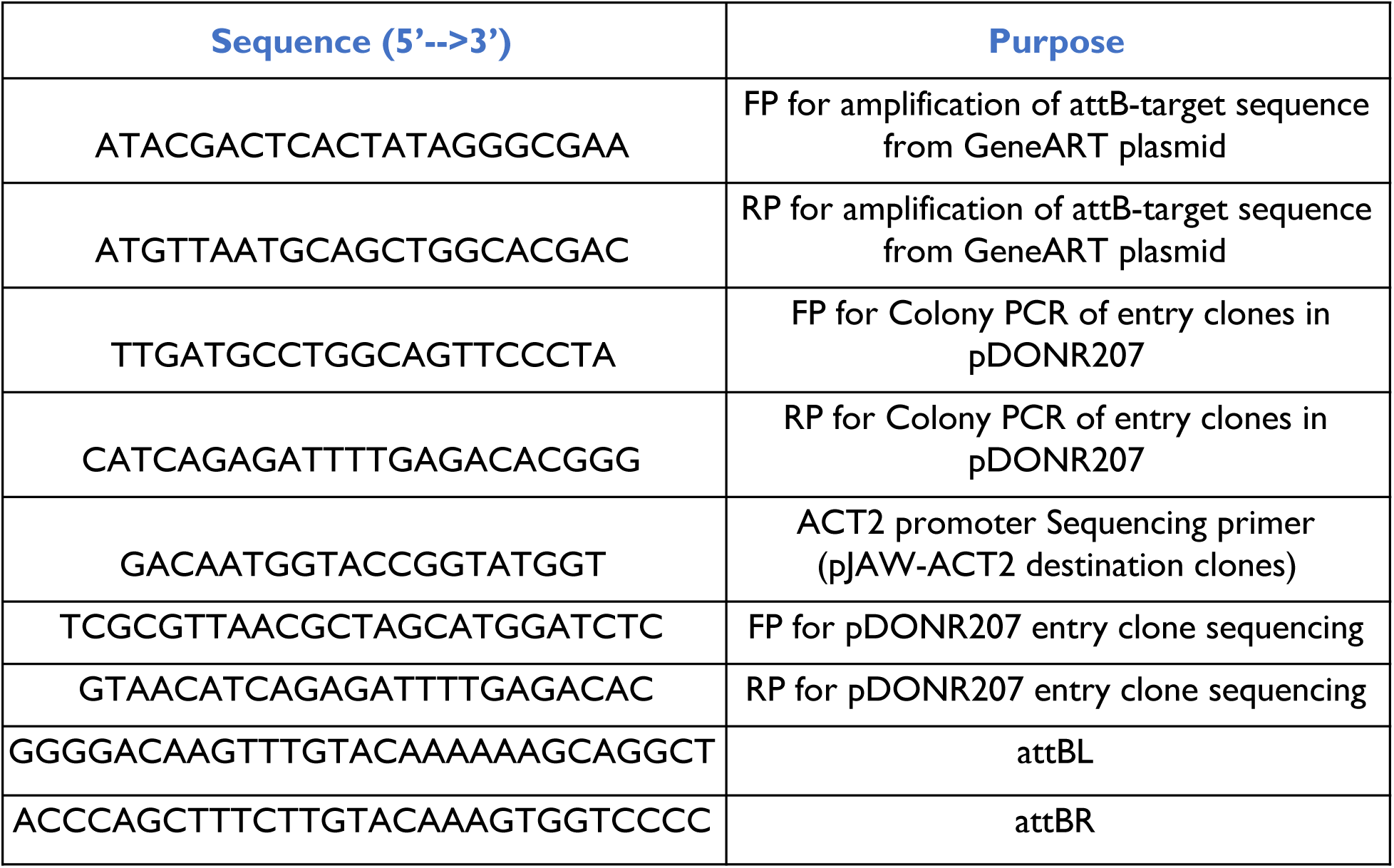
Primer sequences for IR-hairpin cloning (FP forward primer, RP reverse primer)

**Table S2.**
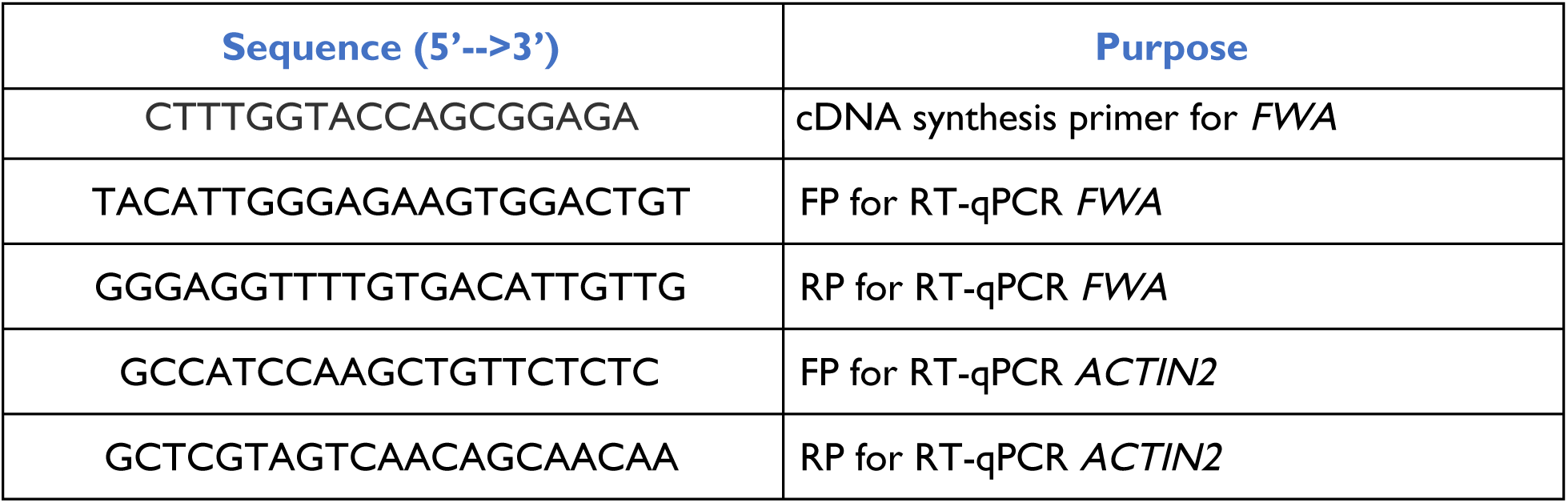
Primer sequence for RT-q-PCR analysis

**Table S3.**
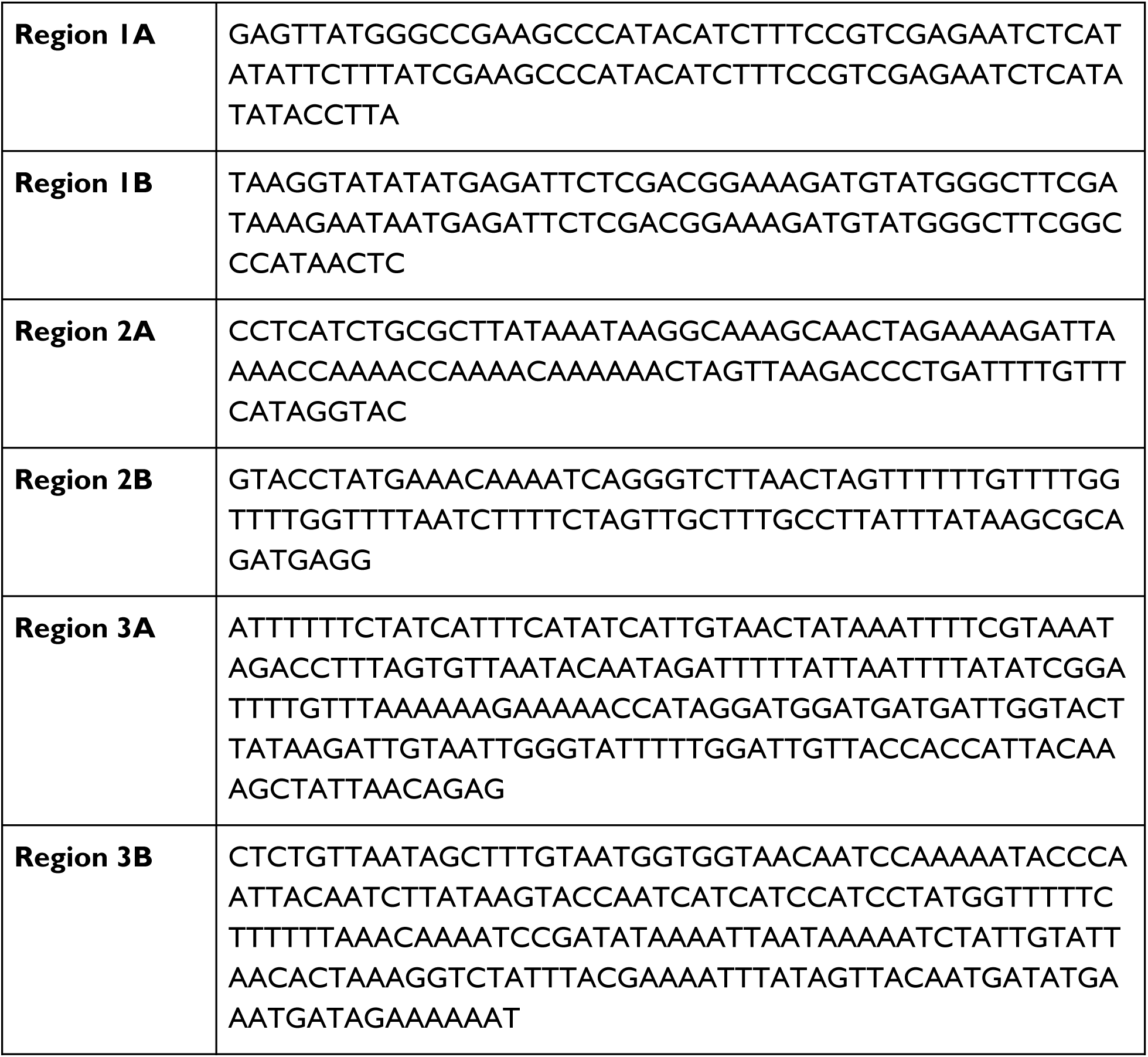
Regions targeted for methylation [Sequences in 5’-->3 orientation]

**Table S4.** Recombinant plasmids generated for this study

| **Plasmid** | **Purpose** |
| --- | --- |
| **pTS001** | pJAW-ACT2 Destination vector targeting methylation in Region 1A |
| **pTS002** | pJAW-ACT2 Destination vector targeting methylation in Region 1B |
| **pTS003** | pJAW-ACT2 Destination vector targeting methylation in Region 2A |
| **pTS004** | pJAW-ACT2 Destination vector targeting methylation in Region 2B |
| **pTS005** | pJAW-ACT2 Destination vector targeting methylation in Region 3A |
| **pTS006** | pJAW-ACT2 Destination vector targeting methylation in Region 3B |

